# Regulating enzymatic reactions in *Escherichia coli* utilizing light-responsive cellular compartments based on liquid-liquid phase separation

**DOI:** 10.1101/2020.11.26.395616

**Authors:** Zikang Huang, Lize Sun, Genzhe Lu, Hongrui Liu, Zihan Zhai, Site Feng, Ji Gao, Chunyu Chen, Chuheng Qing, Meng Fang, Bowen Chen, Jiale Fu, Xuan Wang, Guo-Qiang Chen

## Abstract

Enzymatic reactions in cells are well organized into different compartments, among which protein-based membraneless compartments formed through liquid-liquid phase separation (LLPS) are believed to play important roles^1,2^. Hijacking them for our own purpose has promising applications in metabolic engineering^3^. Yet, it is still hard to precisely and dynamically control target enzymatic reactions in those compartments^4^. To address those problems, we developed Photo-Activated Switch in *E. coli* (PhASE), based on phase separating scaffold proteins and optogenetic tools. In this system, a protein of interest (POI) can be enriched up to 15-fold by LLPS-based compartments from cytosol within only a few seconds once activated by light, and become fully dispersed again within 15 minutes. Furthermore, we explored the potentiality of the LLPS-based compartment in enriching small organic molecules directly via chemical-scaffold interaction. With enzymes and substrates co-localized under light induction, the overall reaction efficiency could be enhanced. Using luciferin and catechol oxidation as model enzymatic reactions, we found that they could accelerate 2.3-fold and 1.6-fold, respectively, when regulated by PhASE. We anticipate our system to be an extension of the synthetic biology toolkit, facilitating rapid recruitment and release of POIs, and reversible regulation of enzymatic reactions.

## Introduction

Up to now, there have been many endeavors contributed to improving the efficiency of manufacturing desired bioproducts in bacteria chassis like *E. coli*. Since simply increasing the expression level of enzymes has many limitations such as adding more burden of protein synthesis to bacteria^5^, poisoning them by deleterious intermediates^6^, etc., finding tools to increase chemical productivity at a fixed enzyme expression level has been a long lasting interest in metabolic engineering. Adjusting the spatial arrangement of enzymes has been proposed as a solution for those requests^6^. With the enzymes in the same pathway assembled together on a DNA or protein-based scaffold, the reaction productivity can be enhanced as the result of minimizing the diffusion distance of intermediates^6,7^. Besides those *de novo* designs, naturally existed bioreactors, known as bacterial microcompartments (BMCs), were hijacked to pack target enzymes of the same pathway together in a protein shell to mitigate intermediate loss^8^. However, the rather large gene set of BMCs^9^, the selectivity of its shell^10^, and the unpredictable efficiency of encapsulation^11^, etc., made it challenging to be reused for customized designs. Furthermore, for all those strategies mentioned, it will be hard to reverse back to the previous state, in case we need to balance different metabolic pathways and bacteria growth regularly^37^. Recently, LLPS-based technologies have been utilized as a much easier method to regulate the spatial distribution of target proteins, since a single protein segment can readily form compartments in cells^1^. By cleaving LLPS domains or soluble tags from a fusion protein, they could be either assembled or disassembled^12^. Other works, taking advantage of optogenetics, could achieve regulating the aggregation of enzymes by light in eukaryotes more dynamically^3,13^. However, it would nevertheless take several to tens of minutes to achieve the final regulatory effect. Moreover, most functional studies of protein spatial engineering *in vivo* focused on regulating multi-step pathways, based on the principle of limiting the diffusion of intermediates^14^, while single-step reaction regulatory tools are still in their early days^15,16^.

Actually, naturally existed membraneless organelles can regulate enzymatic reactions using an alternative strategy: enriching both enzymes and substrates into a limited space. For example, it has been reported that to enrich substrate RNAs directly by interior RNA binding proteins, eukaryotic and prokaryotic organelles can accelerate their processing or degradation^17,18^. Beyond macro-biomolecules, recent studies reported that organic dyes could also be directly incorporated into compartments *in vitro*^19^, providing the potentiality of enriching substrate of single-step reactions by protein-substrate interaction (Figure 1A). Therefore, the overall productivity of an enzymatic reaction can be regulated by switching enzymes and substrates into co-localized or separated states (Figure 1B).

**Figure 1.**
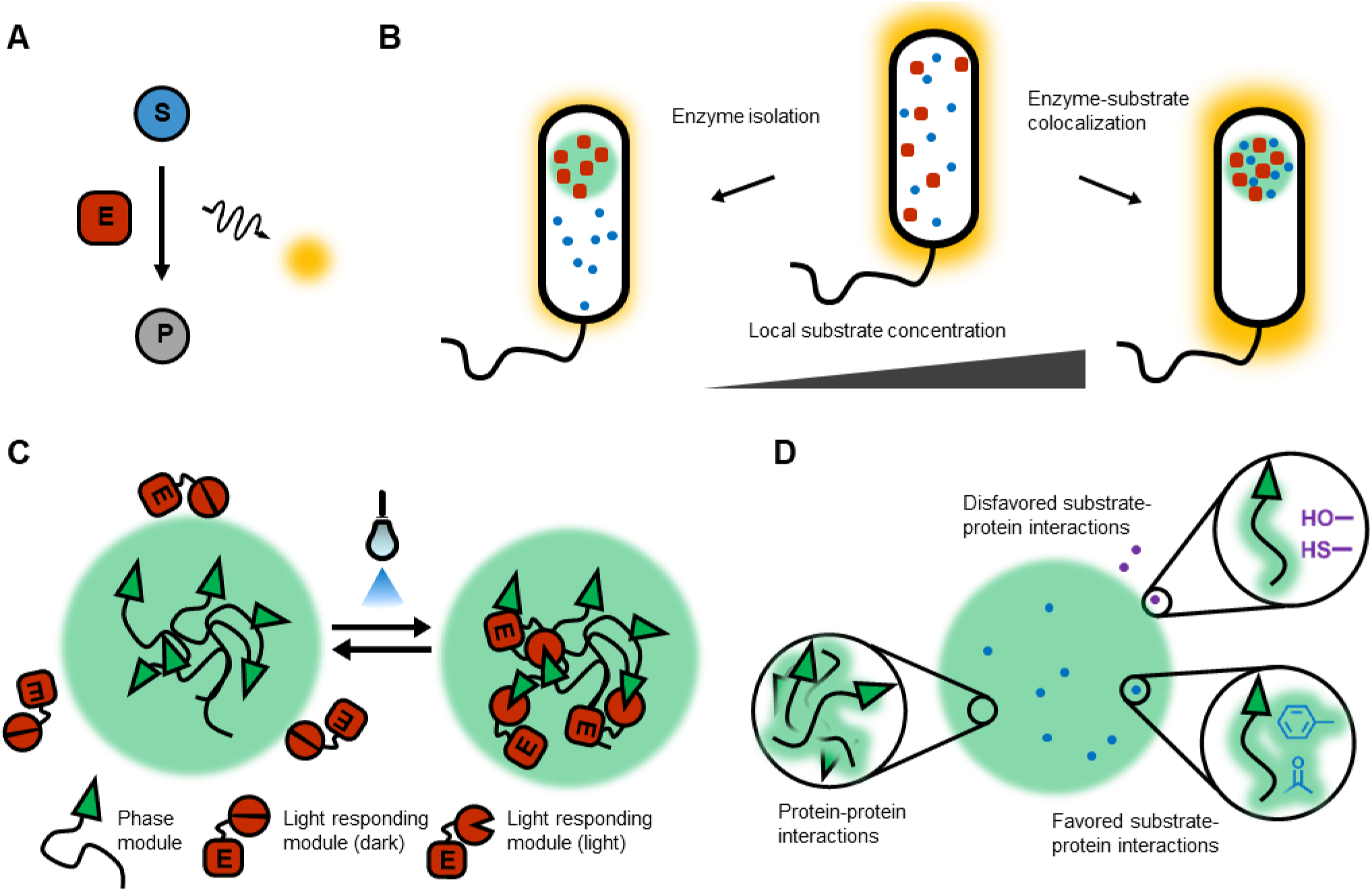
Photo-activated switch for controlling enzymatic reactions. A. Illustration of a model enzymatic reaction. Substrate (S) can be converted to product (P) by enzyme (E), during which light emission by non-stable intermediates directly reflects reaction rate. **B. Enzymatic reaction organized by subcellular compartments.** Under the same overall concentration of enzyme and substrate, a reaction will accelerate if enzyme and substrate co-localize in a limited space, resulting in increasing local substrate concentration, thus increasing effective collision between substrate and enzyme (right). On the other hand, if enzyme is isolated from substrate, the reaction rate will decrease (left). **C. General principle of rearranging enzyme distribution by light.** Each individual of a light responding protein pair are fused with either phase separating protein or enzyme of interest (EOI) to form phase module or light responding module. The phase module can phase separate into protein based compartment. Upon light induction, structural change in light responding protein pair will allow those two modules to bind to each other, and thus EOI can entry the compartment. **D. General principle of concentrating chemicals into the compartment.** More substrates will be enriched in the compartment if substrate-protein interactions are similar to protein-protein interactions promoting phase separation (e.g. pi-pi interaction), while substrates with dissimilar ones will possibly be excluded.

Here, we report a novel tool, photo-activated switch in *E. coli* (PhASE), to dynamically regulate enzymatic reactions (Figure 1C, D). To start with, we introduced various scaffold proteins undergoing LLPS, termed phase module, into *E. coli* to construct artificial compartments. Then, fusing protein of interest (POI) with optogenetic tools, termed light-responding module, we confirmed its reversible recruitment into LLPS-based compartments by light-activated interaction between two modules (Figure 1C). The recruitment process could be completed in a few seconds, which was to our knowledge the fastest method of reorganizing the distribution of proteins *in vivo*. Next, taking the advantage of pi-pi interaction during scaffold protein phase separation^20^, we screened substrates with abundant pi-electrons to be absorbed into the compartment (Figure 1D). With enzymes and substrates enriched in the compartment simultaneously, we found that those reactions could indeed accelerate. In conclusion, we propose PhASE as a new strategy for protein arrangement engineering, which can further regulate enzymatic reactions sensitively and reversibly.

## Results

### Introduction of LLPS-based compartments into E. coli

Above all, since there are few reports of artificially introducing LLPS proteins into prokaryotes^4^, we confirmed their compartment constructing abilities in *E. coli*. The phase module mainly included FUSLCD and truncated GCN4. FUSLCD was known for its ability to phase separate via pi-pi interaction^20^, while GCN4 could further facilitate phase separation through oligomerization^21^ (Figure 2A, B). This fusion protein was used to create LLPS-based membraneless compartments in the following experiment. The whole dynamic formation process of this compartment was recorded in Video S1. We found the compartments could emerge almost anywhere of the peripheral cytosol (Figure 2C) and then move around along the cellular membrane (Figure 2D), much similar to the behavior of liquid droplet. Interestingly, they would finally occupy intracellular space in a regular pattern, much like polar bodies^22^. The pattern was homogeneous across different cells and different phase module expression levels (Figure 2E, Supplementary Figure 1). What was more, probably due to the difference between their LLPS ability, other phase separating proteins, such as Cryptochrome 2 (CRY2), known for its homo-oligomerization after blue light induction^23^, formed compartments at only one pole (Supplementary Figure 2). Then, we tested the fluidity of the compartment formed by the phase module, which showed similar recovering capability as in eukaryotes (Figure 2F)^24^. This property guarantees its potentiality of recruiting proteins and other molecules into it. To be noted, due to the limited space inside a bacterium cell, those compartments were much smaller compared to which formed in eukaryotes and *in vitro*, thus more vulnerable to photo-bleaching. Therefore, the recovery of fluorescence might be the result of protein diffusion from other compartments in the same cell, instead of the periphery of the bleached site (Figure 2F).

**Figure 2.**
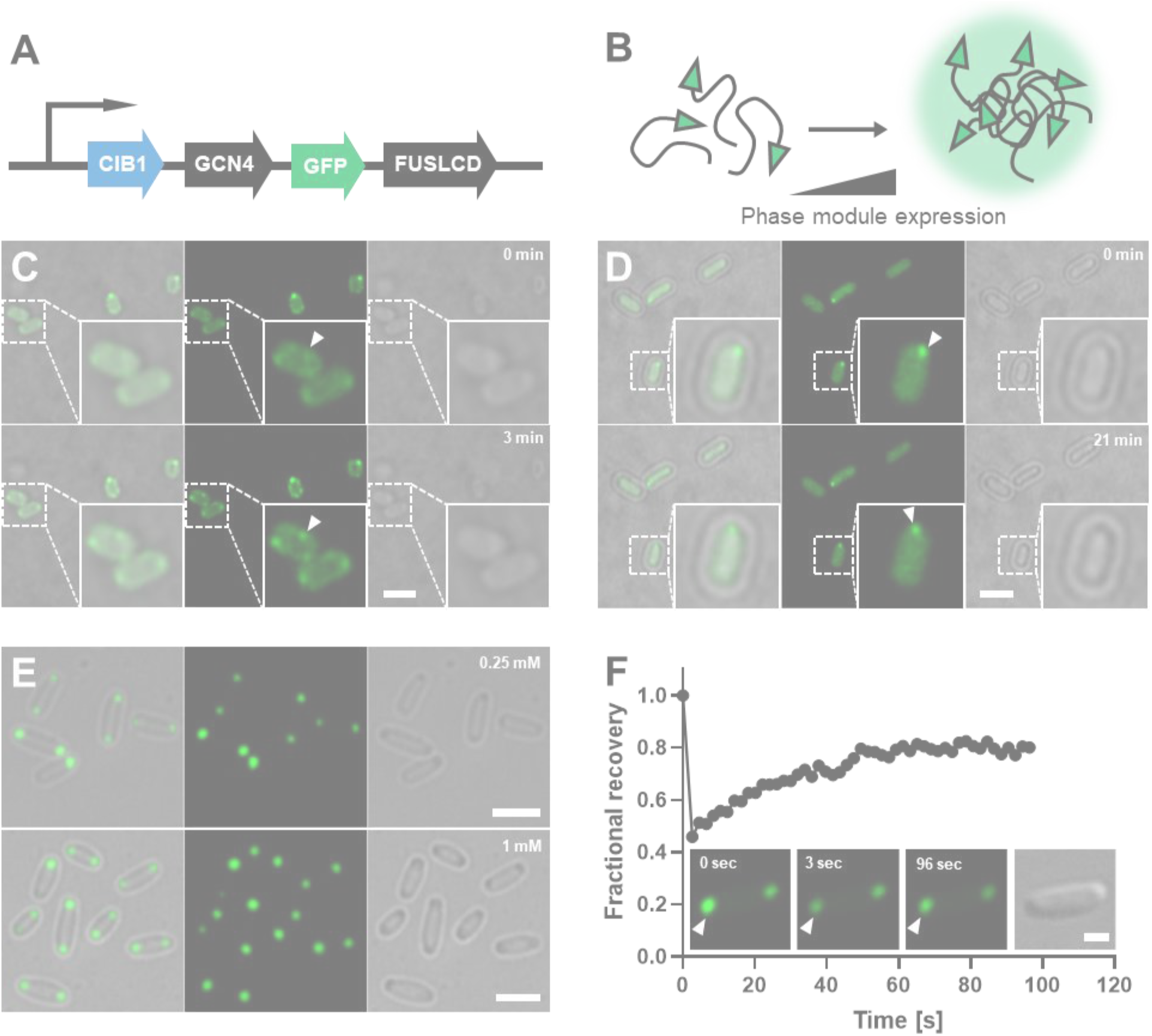
Construction of LLPS based compartment in *E. coli*. A. Illustration of PhASE#1 phase module design. FUSLCD can phase separate *in vivo*, while GCN4 can further facilitate it. CIB1 is one of two individuals of light responding protein pair. See **Figure 3A** for further explanation. **B. Illustration of the mechanism behind compartment construction.** As the concentration of phase module rises beyond a threshold, it will self-aggregate into membraneless compartments, providing a unique chemical environment inside of it. **C, D. The process of compartment formation in *E. coli*.** The nucleation process could happen almost anywhere near the cell wall (C). The compartment then behaves like a droplet, moving around along the cell wall, and finally becomes fixed at both end of the bacteria cell (D). Scale bar, 1 μm. **E. Homogeneous compartment formation in *E. coli*.** Almost every bacterium contains compartments localized at their poles, when expressing PhASE#1 phase module. Phase module was under T7 promoter controlled by *lacO*. Either 0.25 mM or 1 mM IPTG was added to induce its expression. For more IPTG concentration gradients, please refer to Supplementary Figure1. Scale bar, 2.5 μm. **F. Fluidity of PhASE#1 phase module *in vivo*.** Fluorescence Recovery After Photo-bleaching (FRAP) was used to characterize the fluidity of the fusion protein. The recovery of fluorescence might due to rapid protein exchange with other focal planes or cytoplasm. The fluorescence signal of highlighted droplet was normalized to that on the other side of the bacterium. Scale bar, 1 μm.

To further investigate the mechanism behind this special pattern of protein distribution, we transformed phase module into elongated *E. coli* (ΔftsZ) with duplicated genomes but no septum between them, and found protein aggregates formed regularly among genomes (Supplementary Figure 3), similar to what has been reported by Winkler *et, al*^25^, meaning that the curvature of cell wall did not play a role in generating this pattern^26^. What’s more, we found that the compartments formed only along one side of the cell’s major axis, different from where nucleoids aggregated. Combining with the movement of compartments during formation, we reasoned that both LLPS property and the unfavorable contact between nucleoid and phase module may determine its polar localization.

### Reversible recruitment of POI into protein compartments induced by light

To achieve rapid and dynamic recruitment of POI into compartments, we designed the light-responding module to answer the input light signal. Together with the phase module, the system was termed PhASE#1. The phase module acted as a scaffold to construct isolated compartments from the cytosol of *E. coli*, while the light-responding module remained evenly distributed in the cell until triggered by inducing signals. After that, the light-responding module would be recruited into compartments formed by phase module (Figure 1C). Light-responding protein pair CIB1 and CRY2 were chosen to answer the blue light, between which the binding affinity would increase with light stength^27^. CIB1 was fused to phase module, localizing permanently in compartments isolated from cytosol, while CRY2 was fused to the light-responding module, distributing evenly in the cell in darkness. POI was represented by mCherry (Figure 3A). Upon light induction, the interaction between CIB1 and CRY2 will firstly happen at the surface of the compartment, and then, the CIB1-CRY2 complex will move inward gradually and free CIB1 will move outward because of the high fluidity of the compartment. Since protein diffusion is the major time-consuming step in this process, the regulation can be completed in the time scale of seconds theoretically, estimated by protein diffusion rate in *E. coli* given by recent researches^28,29^. Indeed, we found that immediately under 488 nm laser strong stimulation, mCherry could be reorganized into those compartments. The concentration of mCherry inside could reach more than 15-fold compared to the cytosol (Figure 3B) and maintained almost unchanged over 30 minutes (data not shown). The protein compartments labeled with GFP were captured after the stimulation process (Figure 3B), since 488 nm laser was used for light induction of our system. We also tried to stimulate the system with irradiation of lower intensity, the recruited amount of mCherry was reduced, with the concentration enrichment of only about 2-fold (Figure 3C). Furthermore, this recruitment was highly reversible, with the fastest recovery time less than 10 minutes, and a fully reversed state will be attained with no more than 15 minutes (Figure 3C). We next tried to stimulate the system several times after reversion and found that the induction effect was consistent and robust at least in the first three rounds (Figure 3C). The overall reduction of the mCherry signal might be the effect of laser bleaching, but the relative enrichment of the fluorescence signal in the compartment was roughly the same. Therefore, utilizing PhASE# 1, we successfully reorganized POI into a different distribution pattern within seconds in a reversible manner.

**Fig 3.**
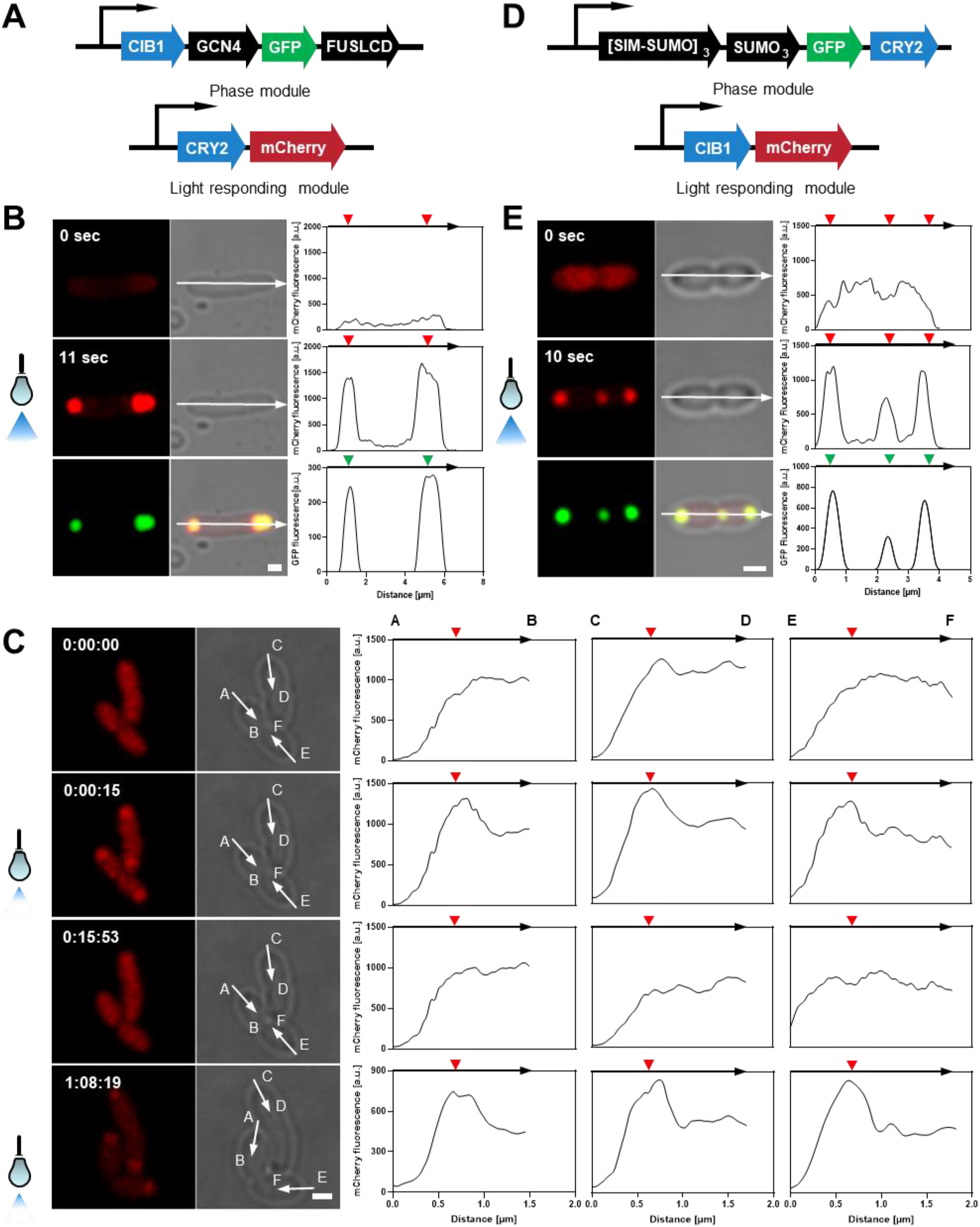
The recruitment of POI into protein compartment. A. Illustration of PhASE#1 system design. CIB1 and CRY2 could bind to each other under 488 nm light. Thus, light responding module would be recruited into protein compartment after light induction. Protein of interest (POI) was represented by mCherry. **B, C. Induction of PhASE#1 system by light of different intensity.** With strong laser irradiation for 8 seconds, the protein compartments could enrich POI for 15-fold compared to cytosol concentration. GFP signal was not shown since its capture also need 488 nm laser, but note that the compartment formed by phase module existed during the whole process. The merged image was captured after the whole stimulation process, thus no time point was labelled. More confirmations of the co-localization of two modules after light were shown in Supplementary Figure 4. (B). With laser irradiation of lower intensity for 3 seconds, the protein compartments enriched POI for 2-fold. After 10 to 15 minutes, the system fully reversed to previous state and was ready for the second round induction (data not shown) and third round of stimulation (1:08:19). With the same set of parameters, this system could be induced as good as the first round. Since the bacteria were actively dividing, the arrow of last time point was not exactly overlapped with that of former time points. The reduction of overall signal intensity was due to the bleaching of mCherry along the manipulation process (C). **D. Illustration of PhASE#2 system design.** Tandem SIM and SUMO, forming multivalent interactions, were used as protein scaffold to construct compartment, while other design principles were the same as PhASE#1. **E. Induction of PhASE#2 system by light.** With strong laser irradiation for 8 seconds, the protein compartments could enrich POI for more than 10-fold compared to cytosol concentration. Scale bar, 1 μm.

We also constructed PhASE#2 system with similar light-responding ability as PhASE#1, substituting FUSLCD with tandem SIM and SUMO repeats (Figure 3D), known for phase separating through multi-valent interactions^30^. The fold enrichment of protein inside the compartment after light induction was over 10 fold (Figure 3E). Besides, since the interaction between phase module and light-responding module in this system can be easily redesigned by adding different number (valency) of SIM or SUMO to light-responding module genetically^30^, we anticipated it as a promising tool for tuning the fold enrichment of POI in the compartment with high temporal precision.

### Regulation of single-step reactions

Besides enzyme concentration, another critical component influencing biochemical reaction rate is substrate concentration. If both enzymes and substrates are congregated in the same compartment, the reaction will accelerate; if they are separated from each other, the reaction will slow down (Figure 1B). Therefore, we set out to explore whether the compartment formed by the phase module can enrich any chemicals. Considering that the main driving force of FUSLCD LLPS is pi-pi interaction^20^, we selected different chemical compounds enriched with pi-electrons, which might be included in the compartment as clients^1^. To explore the regulatory effect of PhASE#1 on single-step reactions, we tethered the enzyme of interest (EOI) to the light-responding module as the experimental group (EXP), so that it could be recruited to the LLPS-based compartment according to light induced CRY2-CIB1 interaction. In the control groups (CTRL), EOIs were fused with mCherry to mimic wild type enzyme and exclude the influence of protein fusion on enzymatic activity (Figure 4A). Then we treated both groups with light and compared the reaction rate difference between them.

**Fig 4.**
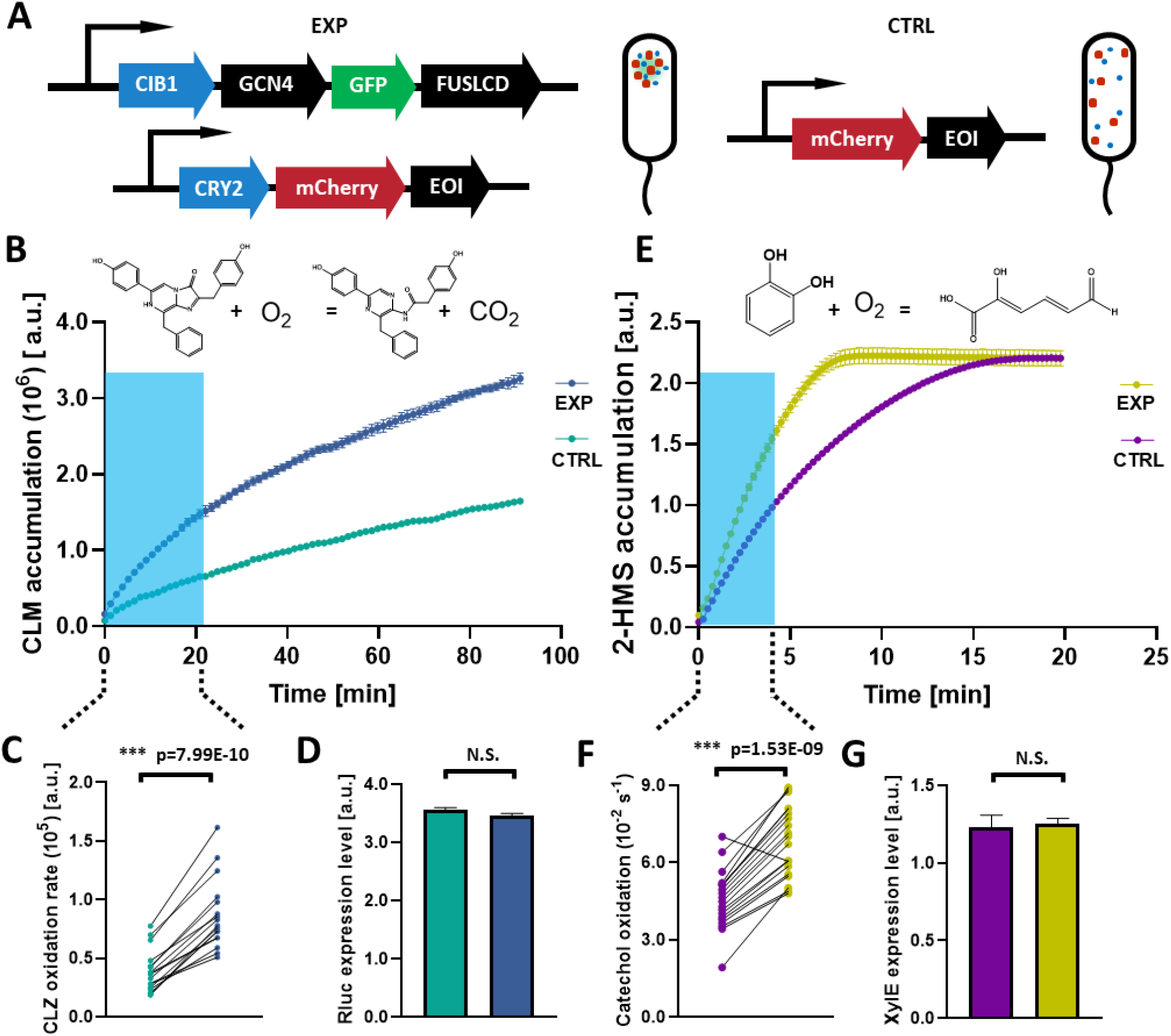
Regulation of single-step enzymatic reactions by PhASE#1 system. A. Module design of PhASE#1 system. In the experimental (EXP) strain, both phase module and light responding module tethered with enzyme of interest (EOI) were transformed, resulting in cells containing a LLPS based compartment that could enrich both enzyme and substrate. In the control (CTRL) strain, only a mCherry fused EOI was transformed, mimicking traditional manufacturing process, with enzyme and substrate dispersed in cells**. B, C, D. Coelenterazine (CLZ) oxidation kinetics.** CLZ can be oxidized by Renilla luciferase (Rluc) (B)^30^. Reaction rate in first 20 minutes from both group, reflected by luminescence at 490 nm, were paired to each other. The experimental group has a significantly higher reaction efficiency (C). Rluc expression level, reflected by mCherry fluorescence, has no significant difference between two groups (D). This experiment has four biological replicates. **E, F, G. Catechol oxidation kinetics.** 1, 2-Dihydroxybenzene (Catechol) can be oxidized to 2-hydroxymuconate semialdehyde (2-HMS) by Catechol 2,3-dioxygenase (XylE) (E)^31^. Reaction rates at first 20 time points were calculated in both groups. The experimental group has a significantly higher reaction efficiency (C). Enzyme expression level, reflected by mCherry fluorescence, has no significant difference between two groups (D). This experiment has four biological replicates.

We first used the coelenterazine (CLZ) oxidation reaction catalyzed by Renilla luciferase (Rluc) as a proof of concept, a reporter system frequently harnessed by *in vivo* studies^31^. The reaction details are shown in Figure 4B^32^, and the luminescence during the process was used as a direct measurement of reaction rate (Figure 4C)^33^. After light treatment, EOIs were successfully recruited into the LLPS-based compartments (Supplementary Figure 4A). In the first 20 minutes after CLZ addition, the EXP group displayed a significantly higher oxidation rate compared to the CTRL group, with 2.6 times of coelenteramide (CLM) accumulation difference. Since the overall substrate and enzyme concentration remained the same among those groups (Figure 4D), we deduced that CLZ should have interacted with the protein scaffold so that it could be absorbed by the compartment from the cytosol, otherwise the limitation of enzyme movement in cells would only harm the reaction efficiency. On the other hand, from the bottom-up perspective, this enrichment was reasonable for the great amount of pi-electrons CLZ contained, making it a potential client for LLPS-based compartment formed through pi-pi interaction between FUSLCD^20^. To be noted, the concentration of bacteria might not have a major impact on this reaction at the proper range (Supplementary Figure 5). With four biological replicates, we concluded that PhASE#1 could indeed regulate this single-step enzymatic reaction.

Next, we set out to try more biochemical reactions with practical use. Catechol 2,3-dioxygenase (XylE) catalyzes the oxidation of catechol, which is a key metabolite in many organic degradation pathways^34,35^ (Figure 4E). Similar to the design above, a modified light-responding module tethered with XylE was constructed and could be successfully enriched into the LLPS-based compartments (Supplementary Figure 4B). Both incubated under daylight, the EXP group showed a much higher reaction rate compared to the CTRL group at the same enzyme and substrate level, with 1.6 times 2-hydroxymuconate semialdehyde (2-HMS) accumulation difference 4.5 minutes after the substrate addition. Unlike CLZ, catechol is a hydrophilic chemical. Thus, though only limited reactions have been tested, our result suggests that hydrophobicity might not be a major driving force of substrate enrichment, while whether a chemical can act as a client of the compartment to interact with phase module really matters.

## Discussion

To tune metabolic reactions by enzyme expression level has been studied quite comprehensively, but there are times when higher protein concentration might be deleterious or even impossible^5,36^. Here, we present a tool that can add another layer of regulation at the post-translational level to further increase reaction efficiency. Moreover, its great reversibility allows the regulation to be dynamical, compared to chemical inducers that cannot be easily removed and scaffolding methods in assembling enzymes that cannot be released^6,7^. To further increase the temporal precision of this tool, we designed a fixed compartment during the whole regulatory period, bypassing the time-consuming compartment assembly process^3^. What’s more, taking advantage of the rather narrow space in *E. coli*, compared to eukaryotic cells, the recruitment will be completed even faster. With this regulatory system, combining with our exploration of chemical interactions between compartment client and scaffold, we expanded the ability of protein aggregates in metabolic regulation of single-step enzymatic reactions.

However, there are still some considerations about this tool. First of all, though using proper phase separating proteins to enrich both target substrate and enzyme in the same compartment could be achieved theoretically, it might be hard to find such scaffolds for every specific chemical in practical^14^. To be noted, too specific interactions between chemical client and protein scaffold might not increase reaction efficiency either, since it might compete substrate binding with enzymes. Secondly, the protein compartment occupies a large amount of space in a prokaryotic cell, which might have impact on bacterial metabolism. Finally, because the substrates are enriched via non-specific interactions, there will still be some of them left outside the compartment, which might result in a high background reaction rate before light induction. Nevertheless, with a very high regulatory temporal resolution and great reversibility, our systems may tune metabolic flux more dynamically, finding a better balance between different pathways and bacteria growth, which has been proved to enhance chemical production^37^. Besides metabolic regulation, our tools also have the potentiality to regulate and study various life activities. By quickly isolating proteins from their substrates or partners, we anticipate it suitable for studying transient biochemical reactions and the switch of signaling pathways (Figure 2)^38^. Some of those processes only last for a few seconds, which can hardly be probed by transcriptional, translational control, or degradation induction approaches such as RNAi, achieving the regulatory effect in minutes or even hours^30,38^. Also, thanks to the high spatial resolution of laser, it is well suited for single-cell manipulation^38^. Another possible application is the inducible bacterial cell differentiation. Expressing CRY2 tethered POI, we may control its asymmetric aggregation and thus asymmetric division by light^3^ (Supplementary Figure 2). The resulting bacteria community will be heterogeneous in POI expression level and aggregation state^25,39^.

In conclusion, we enlarged the toolbox of metabolic engineering in prokaryotes by combining engineered phase separating proteins and optogenetic tools. With PhASE#1 and #2 systems, we can complete the protein redistribution process within only a few seconds and achieve different regulatory states by controlling light intensity. Using CLZ and catechol oxidation reactions as model systems, we successfully filled the gap of regulating single-step reactions by artificially constructed compartments *in vivo* and raised a potential principle in designing such tools based on chemical interaction between compartment scaffold and client. We anticipated this tool to facilitate precise metabolic engineering as well as broad life activity manipulation.

## Methods

### DNA manipulation

All constructs were assembled using available restriction enzymes and T4 ligase, or Hi-Fi assembly from New England BioLabs (NEB) and other basic molecular cloning approaches. To construct the light-responding LLPS system PhASE#1, we acquired gene sequences of FUSLCD (FUS residues 1-212, NP_004951.1), CRY2 (NP_171935.1), CIB1 (NP_195179.2), and GCN4 (residues 2-34, 1W5I_A) from Pilong Li’s group at Tsinghua University. The sequences of the other system, PhASE#2, contained SIM (E3 SUMO-protein ligase PIAS2 isoform X1 residues 505-527, XP_006722634.1) and SUMO (XP_012635485.1), were synthesized by Ruibiotech directly. Since we only used CRY2 as a light-responding element, we added an MBP tag to avoid its aggregation^40^. Plasmids contain genes encoding Rluc (Renilla luciferase, AGU01696.1), XylE (catechol 2,3-dioxygenase, WP_011005909.1), fluorescence proteins were acquired from iGEM parts distribution. All sequence ID could be found on NCBI.

### Plasmid transformation

For plasmid amplification, we transformed 1 μL plasmid (about 100 ng μL^−1^) to 50 μL *E. coli* DH5 chemical competent strain. It was then incubated on ice for 25 minutes followed by 90 seconds heat shock. Then it would be mixed with 450 μL LB medium and shook for an hour. For protein expression, we utilized *E. coli* BL21(DE3) strain. We used 100 μL for co-transformation of two plasmids and added 900 μL LB medium to activated. For single plasmid transformation, the protocol was the same as *E. coli* DH5□ amplification. For plasmid construction, often by HiFi assembly (NEB), we transformed 5 μL product to 100 μL bacteria solution and followed by the same protocol as amplification.

### Strains, media, and culture conditions

LB medium and plates were used for *E. coli* cultivation. Kanamycin and ampicillin were added at 100 mg L^−1^ when used alone, 50 mg L^−1^ when combined. For plasmid amplification and molecular cloning, we used *E. coli* DH5□, cultivated at 37°C, 220 r.p.m. For protein expression experiments, we used *E. coli* BL21(DE3) strains (TIANGEN and TSINGKE) and regulate protein expression level by lacO, adding different concentrations of IPTG. Seed cultures were grown at 37°C, 220 r.p.m, and adjusted to OD600 of 0.6-1.0 for IPTG induction. Since phase separating proteins were known to form inclusion bodies at high expression levels and may cause problematic folding when fused to other proteins, we express all fusion proteins at 16°C, 220 r.p.m^40^, while test light response and enzymatic reactivity at room temperature or appropriate temperatures.

### Confocal microscopy

All images shown in this paper were captured by Nikon A1 LFOVT. To assay bacteria growth (Figure 2C, D) and light response (Figure 3B, C, E) for a long period of time, we used an LB agar pad to fix 1 μL overnight *E. coli* under a microscope. The pad layer was solid 1% LB agarose, while 400 μL was added to each glass coverslip-bottomed (20 mm in diameter) 35 mm Petri dish (D35-20-1-N). For confirmation of bacteria morphology and compartment localization pattern (Supplementary Figure 2–4), recruitment of enzymes into compartments (not shown), and other short period observation, we used glass slides to fix them. All images captured were used a 100X oil immersed lens. 488 nm laser was used only during observation of phase separation since light-responding proteins were sensitive to its induction. Therefore, 561 nm laser was the only light source at light induction experiments. ProLong Live Antifade Reagent (P36975) was used to avoid photo-bleaching of mCherry (1:50 dilution). Strong light induction was performed using region of interest (ROI) to guide 488 nm lasers, 3% lasted for 8 seconds in Figure 3B and E, while weak light induction was conducted with 2% laser intensity for 6 seconds. FRAP assays were carried out using slightly higher laser power with ROI as a single pixel.

### Image analysis

All confocal data analyses were performed on NIS Element Analysis. All parameters in the same time series were kept the same. All quantified fluorescence intensity was calculated before images were smoothed for display usage. The contrast of images was enhanced by adjusting LUT.

### Live staining of bacteria DNA

We utilized NucBlue Live Cell Stain ReadyProbes reagent (R37605) to label *E. coli* nucleoid and LLPS based compartment. 2 drops of the solution were added to 1 ml bacteria culture and incubated in darkness for 15 minutes. The observation should be done in 45 minutes.

### Coelenterazine oxidation assay

CTRL and EXP group bacteria were collected by centrifugation of 5,000-8,000 r.p.m, 2 minutes, and washed two times by PBS. Then, 100 μl PBS culture was added to each well of the 96-well plate, and the mCherry fluorescence level of EXP groups and CTRL groups were matched to each other. Prior to each assay, 10 mM CLZ ethanol solution was diluted in PBS to a final concentration of 250 μM and incubated for 45 minutes to an hour. During the experiment, 3 μl CLZ diluted solution was added to each well^31^. The luminescence of 490 nm was measured at 21-27°C. CLZ was bought from Invitrogen (C2944). Thermo Scientific Varioskan LUX was used for quantitating reactions. SkanIt software was used for data management.

### Catechol oxidation assay

CTRL and EXP group bacteria were collected by centrifugation of 5,000-8,000 r.p.m, 2 minutes, and washed two times by PBS. Then, 100 μl PBS culture was added to each well of 96-well plate, and the mCherry fluorescence level of EXP groups and CTRL groups were matched to each other. Prior to each assay, 100 mM catechol water solution was diluted to 10 mM working solution, 2.5 μL of which was then added to each well. The absorbance of oxidation product 2-HMS at 377 nm can be measured at 19-22°C. Catechol was bought from damas-beta (120-80-9). Thermo Scientific Varioskan LUX was used for quantitating reactions. SkanIt software was used for data management.

## Acknowledgement

We thank members of Pilong Li’s lab for helpful discussions and kindly provision of plasmids. We also thank Dr. Peng Li for his team and lab training and members of iGEM Tsinghua team (2018) for their guidance in brainstorming. This work was supported by Tsinghua University Initiative Scientific Research Program, Tsinghua Xuetang Program, and Student Research Training Program.

## Supplementary figures

**Supplementary Figure 1.**
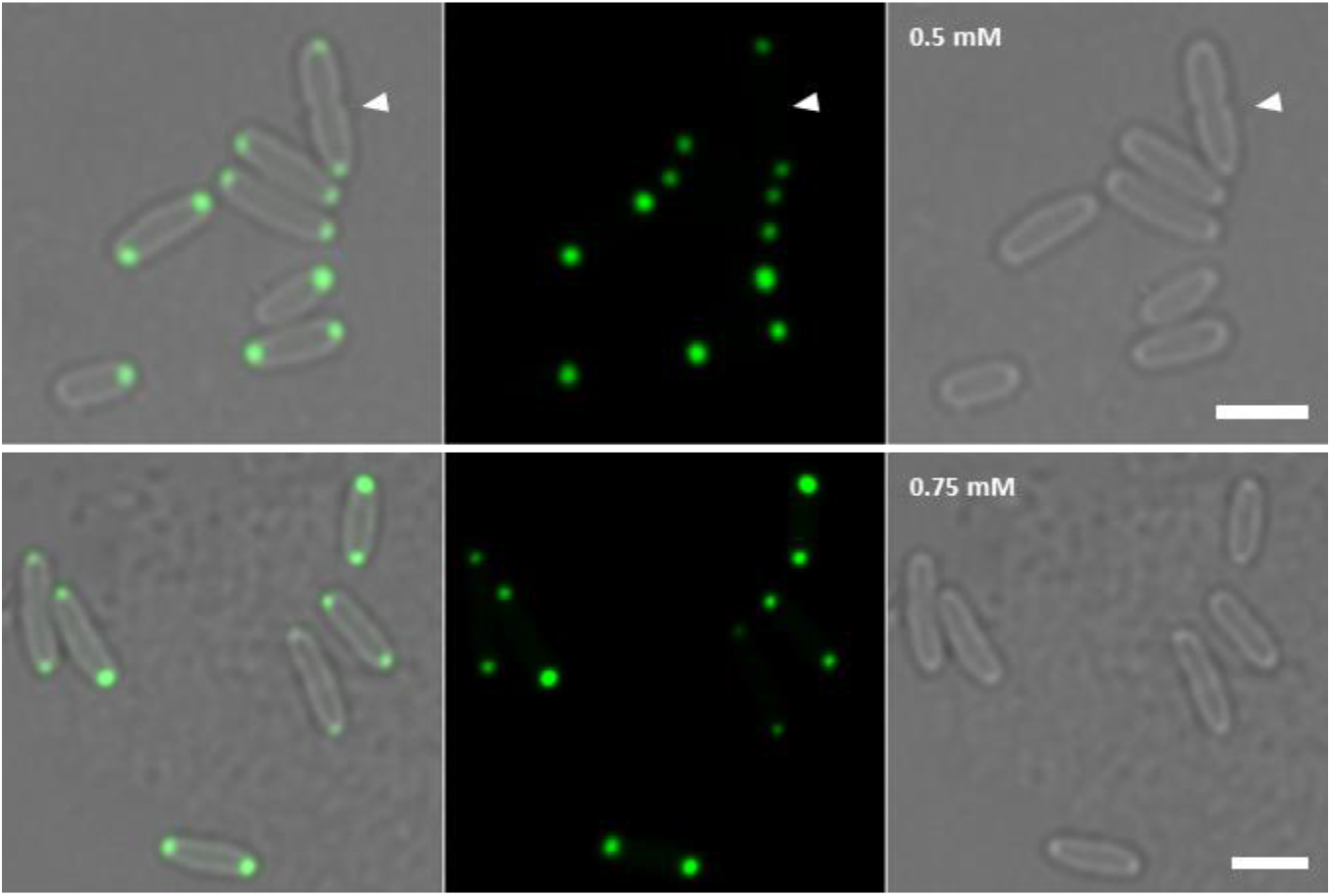
Homogeneous compartment formation by phase module. Almost all *E. coli* cells expressing phase module have similar compartments localized at both poles of bacteria. Phase module was under T7 promoter controlled by *lacO*. Either 0.5 mM or 0.75 mM IPTG was added to induce its expression. To be noted, those two compartments were often evenly inherited by daughter bacteria after dividing (white arrow). Therefore, some short bacteria may only contain one compartment. Scale bar, 2.5 μm.

**Supplementary Figure 2.**
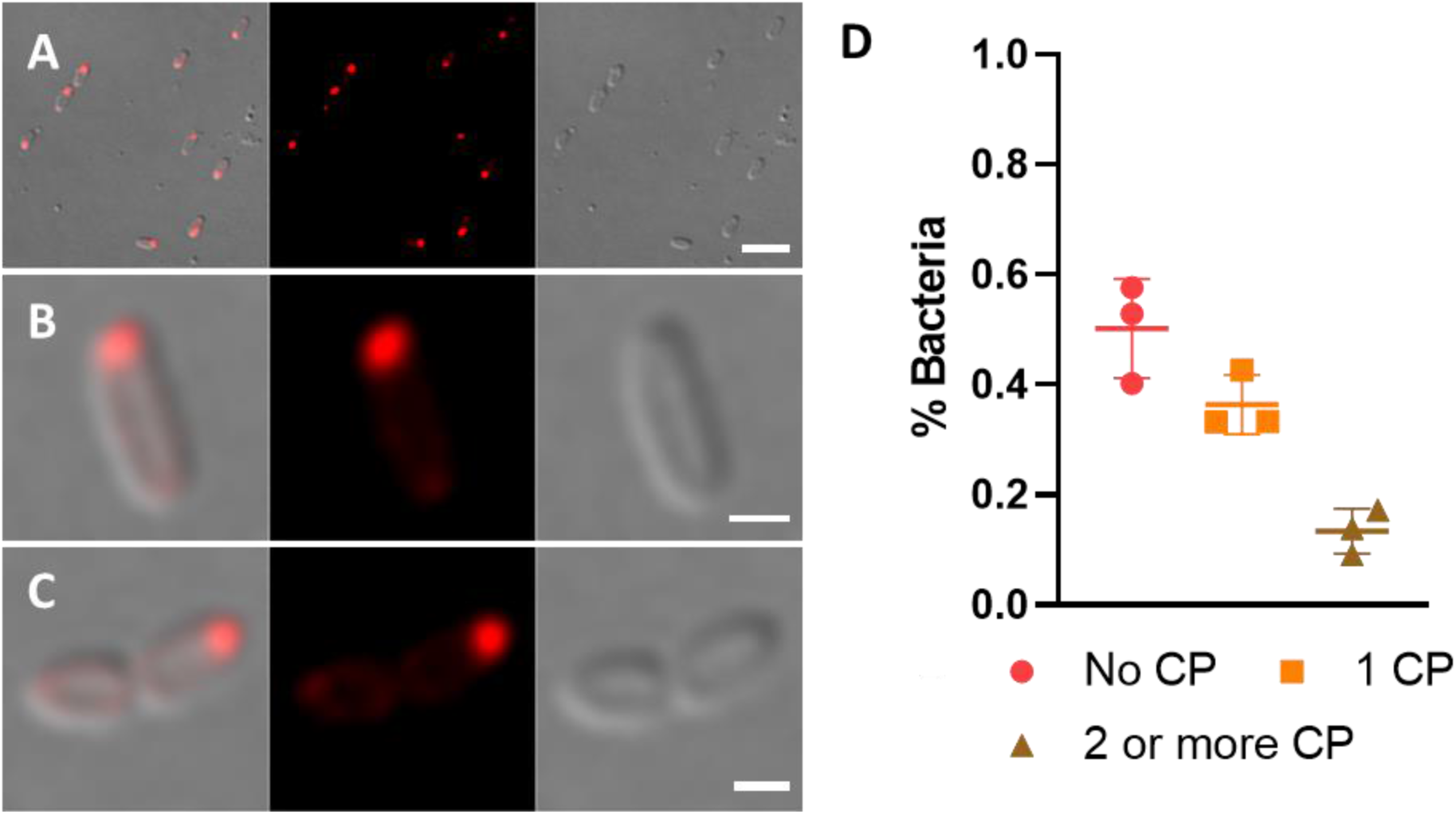
Various scaffold proteins construct compartments with different distribution patterns. **A.** Almost all *E. coli* cells expressing CRY2-mCherry only form a single aggregate at one end of the cell. Scale bar, 5 μm. **B.** Detailed illustration of uni-polar compartment localization. **C.** The uni-polar localized compartment may lead to asymmetric inheritance of protein level in daughter cells. B, C scale bar 1 μm. **D.** Statistics of compartment (CP) number in a bacterium. All fields counted have more than 200 bacteria.

**Supplementary Figure 3.**
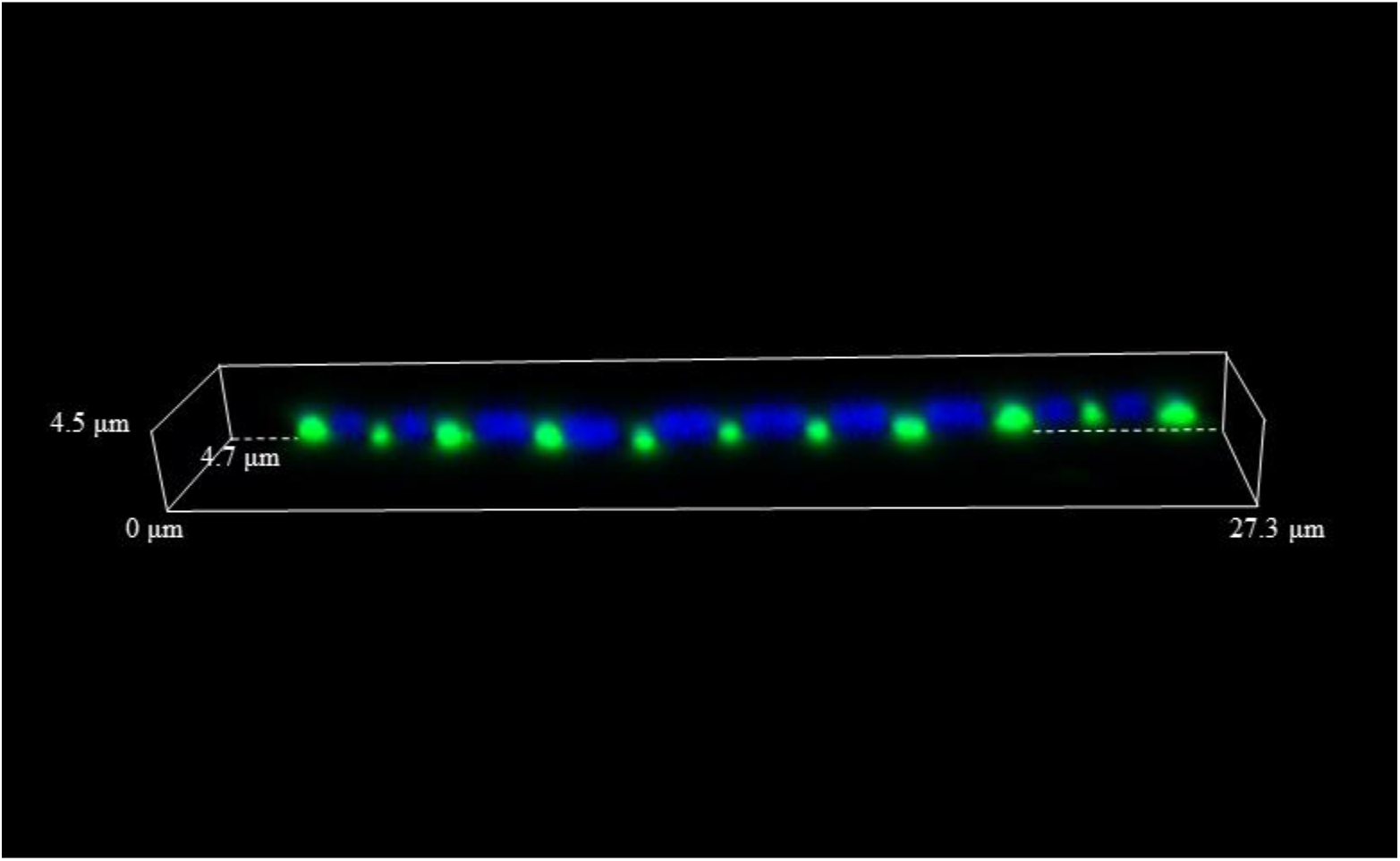
Nucleoid exclusion might be a potential driving force of polar localization of protein based compartment. **A.** Z-stack reconstitution of elongated *E.coli* (ΔftsZ) with several nucleoids in one cell but no septums. It displayed an interesting pattern with compartments formed among nucleoids. To be noted, they form only along one side of the cell’s major axis, while nucleoids aggregate along the other side. This might be another evidence supporting protein droplets prefer occupying space where the contact with nucleoids can be minimized.

**Supplementary Figure 4.**
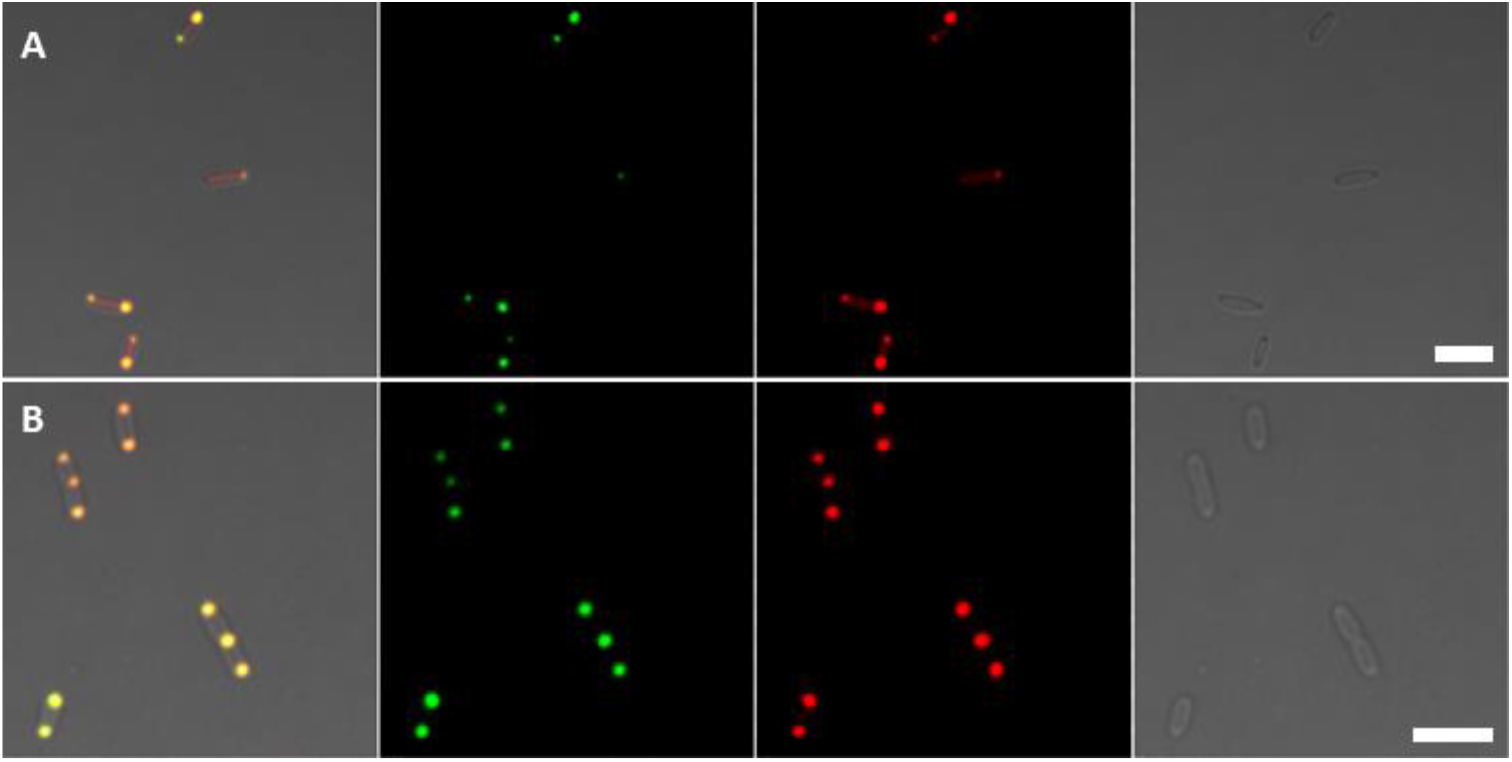
Recruitment of EOIs into LLPS-based compartments by PhASE#1 system after photo-activation. A, B. Rluc and XylE were used as EOI, respectively. They were enriched in the compartments successfully. Scale bar, 5 μm.

**Supplementary Figure 5.**
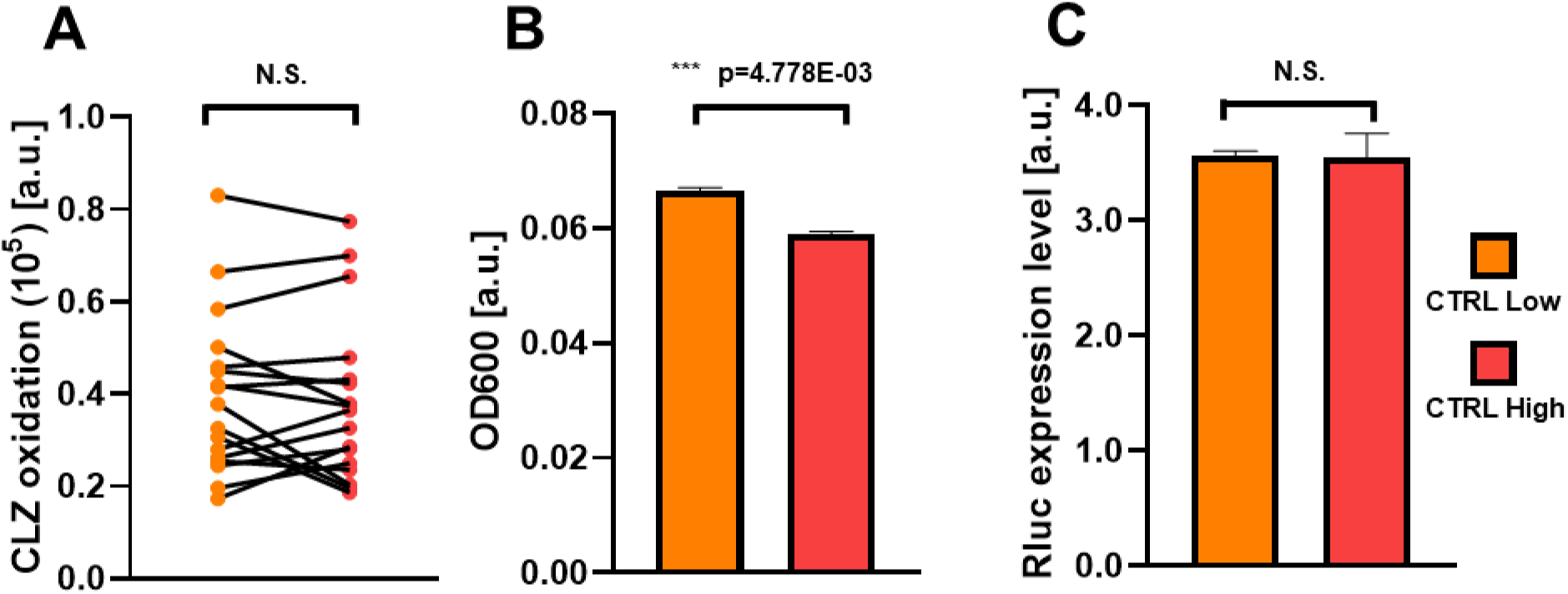
CLZ oxidation rate difference between EXP and CTRL was not due to different bacteria concentration. **A.** There was no significant difference in CLZ oxidation rate between two CTRL strains with different enzyme expression level. **B, C.** The enzyme concentration was adjusted to the same level, while bacteria concentration has a significant difference between two groups. CTRL Low and High indicated enzyme expression induced by 0.02 mM or 0.5 mM IPTG under same culturing condition.

